# A shared *cis*-regulatory module activates transcription in the suspensor of plant embryos

**DOI:** 10.1101/297309

**Authors:** Kelli F. Henry, Anhthu Q. Bui, Tomokazu Kawashima, Robert B. Goldberg

## Abstract

The mechanisms controlling the transcription of gene sets in specific regions of a plant embryo shortly after fertilization remain unknown. Previously, we showed that G564 mRNA, encoding a protein of unknown function, accumulates to high levels in the giant suspensor of both Scarlet Runner Bean (SRB) and Common Bean embryos, and a *cis*-regulatory module containing three unique DNA sequences, designated as the 10-bp, Region 2, and Fifth motifs, is required for *G564* suspensor-specific transcription [Henry, K. F. *et al., Plant Mol. Biol*. 88(3):207-217 (2015); Kawashima, T. *et al., Proc. Natl. Acad. Sci USA* 106(9):3627-3632 (2009)]. We tested the hypothesis that these motifs are also required for transcription of the SRB *GA 20-oxidase* gene, which encodes a gibberellic acid hormone biosynthesis enzyme and is co-expressed with *G564* at a high level in giant bean suspensors. We used deletion and gain-of-function experiments in transgenic tobacco embryos to show that two *GA 20-oxidase* DNA regions are required for suspensor-specific transcription – one in the 5’ untranslated region (UTR) (+119 to +205) and another in the 5’ upstream region (−341 to −316). Mutagenesis of sequences in these two regions determined that the *cis*-regulatory motifs required for *G564* suspensor transcription are also required for *GA 20-oxidase* transcription within the suspensor, although the motif arrangement differs. Our results demonstrate the flexibility of motif positioning within a *cis*-regulatory module that activates gene transcription within giant bean suspensors, and suggest that *G564* and *GA 20-oxidase* comprise part of a suspensor gene regulatory network.

**Significance:** Little is known about how genes are expressed in different plant embryo regions. We tested the hypothesis that shared *cis*-regulatory motifs control the transcription of genes specifically in the suspensor. We carried out functional studies with the Scarlet Runner Bean (SRB) *GA 20-oxidase* gene that encodes a gibberellic acid (GA) hormone biosynthesis enzyme, and is expressed specifically within the suspensor. We show that *cis*-regulatory motifs required for *GA 20-oxidase* transcription within the suspensor are the same as those required for suspensor-specific transcription of the SRB *G564* gene, although motif number, spacing and order differ. These *cis*-elements constitute a control module that is required to activate genes in the SRB suspensor and may form part of a suspensor regulatory network.

## Introduction

In most higher plants, embryogenesis begins with the asymmetric division of the zygote to give rise to a small apical cell and a large basal cell (1). The apical and basal cells follow distinct pathways to differentiate into an embryo proper and suspensor, respectively (2, 3). Whereas the embryo proper undergoes many developmental and morphological changes to eventually become the mature embryo within the seed, the suspensor is a terminally differentiated embryo region that degenerates as the embryo matures. Several studies have shown that different genes are expressed in the embryo proper and suspensor (4-8), but how these genes are organized into regulatory networks (9) operating in the different embryo regions remains unknown.

Previously, we began to dissect the gene regulatory networks programming early embryo development by analyzing the activation of *G564*, a gene encoding a protein of unknown function that is active specifically in the giant suspensors of Scarlet Runner Bean (*Phaseolus coccineus*; SRB) and Common Bean (*Phaseolus vulgaris*) (Fig. 1*A-E*) (10-12), which diverged ~2 million years ago (13). *G564* suspensor transcription is activated by five motifs: (i) three 10-bp motifs with the consensus 5’-GAAAAGCGAA-3’ that can tolerate up to three non-adjacent mismatches, (ii) a Region 2 motif 5’-TTG(A/G)(A/G/T)AAT-3’ and (iii) a Fifth motif 5’-(A/G)AGTTA-3’ (Fig. 2) (11, 14).

**Fig. 1.**
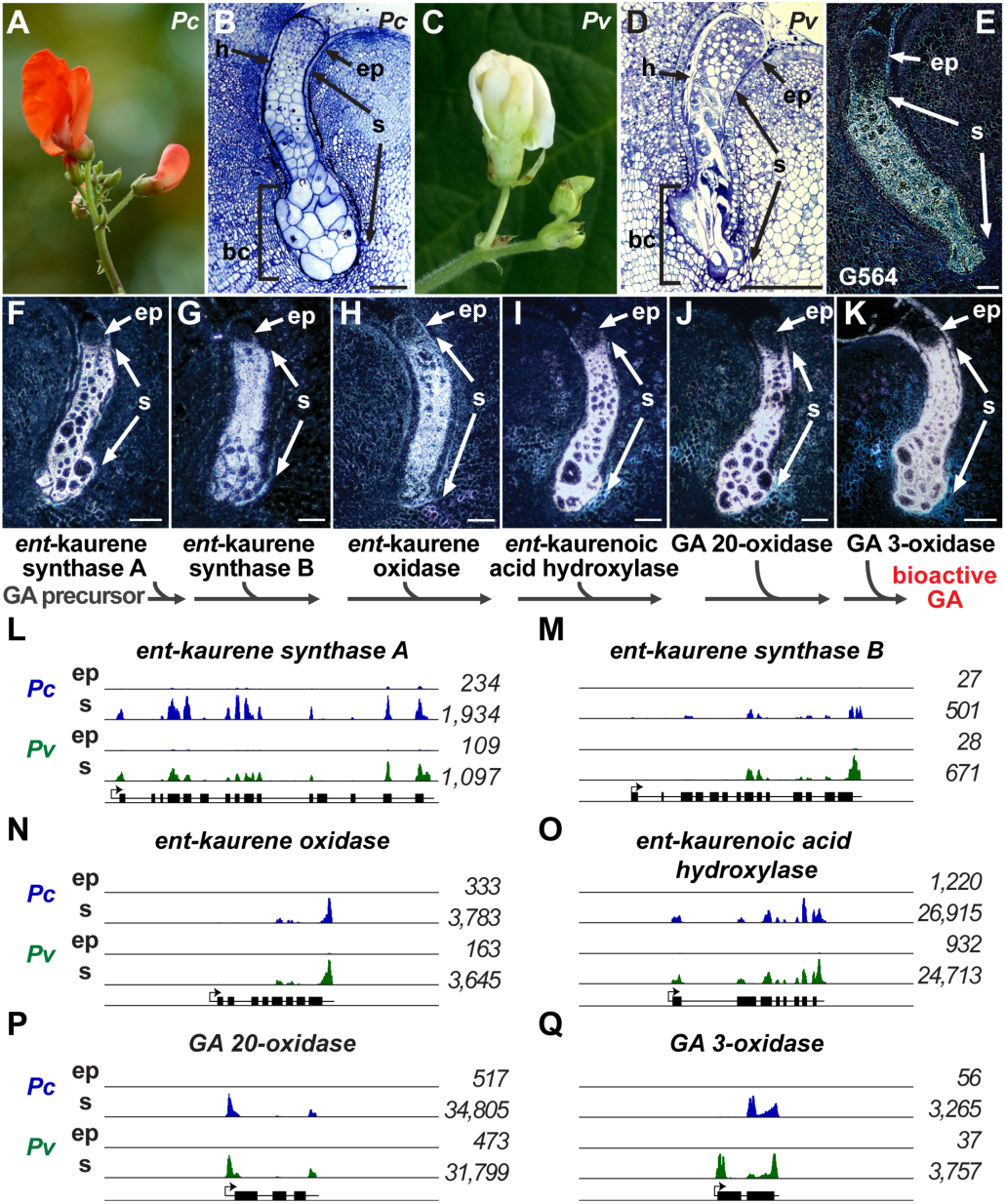
Localization of GA biosynthesis pathway enzyme mRNAs in SRB and Common Bean embryos. (*A*) SRB flower. (*B*) Plastic section of SRB globular-stage embryo taken from Henry and Goldberg (2015) (10). (*C*) Common Bean flower. (*D*) Paraffin section of Common Bean globular-stage embryo taken from Henry and Goldberg (2015) (10). (*E*) Localization of G564 mRNA in a SRB globular-stage embryo taken from Weterings *et al*. (2001) (12). (*F-K*) *In situ* localization of GA biosynthesis enzyme mRNAs in globular-stage SRB seeds: ent-kaurene synthase A (*F*), ent-kaurene synthase B (*G*), *ent*-*kaurene oxidase* (*H*), ent-kaurenoic acid hydroxylase (*I*), GA 20-oxidase (*J*), GA 3-oxidase (*K*). Photographs were taken using dark-field microscopy. (*L-Q*) Genome browser views of RNA-Seq coverage of: *ent*-kaurene synthase A (*L*), *ent*-kaurene synthase B (*M*), *ent*-kaurene oxidase (*N*), *ent*-kaurenoic acid hydroxylase (*O*), GA 20-oxidase (*P*), and GA 3-oxidase (*Q*) in SRB and Common Bean globular-stage suspensor and embryo proper regions. RNA-Seq data were taken from GEO accession GSE57537. Numbers indicate average reads per kilobase per million of two biological replicates. Each panel depicts an 8-kb window including the gene structure. Black boxes represent exons. Black lines represent UTRs and introns. Arrows indicate the transcription start site. bc, basal cells; ep, embryo proper; h, hypophysis region; *Pc*, *P. coccineus*; *Pv*, *P. vulgaris*; s, suspensor. (Scale bar: 50 µm.)

**Fig. 2.**
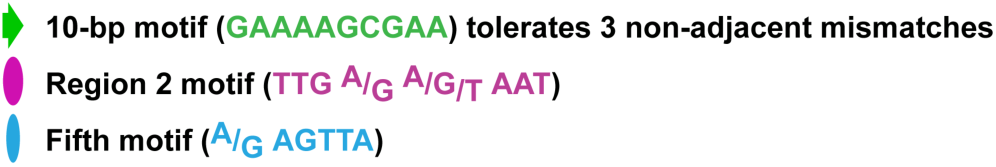
Consensus sequences for suspensor 10-bp motif, Region 2 motif and Fifth motif. Consensus sequences were generated from *G564* DNA sequences shown to be required for transcription within the suspensor (11, 14).

In this paper we test the hypothesis that genes with similar suspensor-specific expression patterns in giant bean suspensors utilize a shared *cis*-regulatory module (9) with common *cis*-control elements. We show that genes encoding enzymes for each step of the gibberellic acid (GA) biosynthesis pathway (15) are expressed at high levels in SRB and Common Bean globular-stage suspensors, similar to *G564*, suggesting that these genes are co-regulated. We analyzed in detail the upstream region of one gene in the GA pathway, SRB *GA 20-oxidase*, and present experiments demonstrating that the *GA 20-oxidase* upstream region can activate suspensor transcription in globular-stage tobacco embryos. Deletion, gain-of-function (GOF), and mutation analyses in transgenic tobacco embryos showed that the *GA 20-oxidase* upstream region −341 to +238 is sufficient for suspensor-specific transcription, and contains functional *cis*-regulatory elements that are also required for suspensor transcription of the SRB *G564* gene (11, 14). Mutagenesis of the predicted suspensor *cis*-regulatory elements in the *GA 20-oxidase* upstream region showed that sequences similar to the 10-bp motif, Region 2 motif, and Fifth motif are required for *GA 20-oxidase* suspensor transcription. Our results demonstrate that transcription of the *G564* and *GA 20-oxidase* genes within the SRB suspensor is activated using a shared *cis*-regulatory module that differs in the number, spacing and order of *cis*-motifs, and that this *cis*-regulatory module may form part of a gene regulatory network that operates in giant bean suspensors shortly after fertilization.

## Results

### mRNAs encoding GA biosynthesis enzymes localize to the SRB and Common Bean giant suspensor

We carried out *in situ* hybridization analysis on SRB globular-stage seeds to determine the mRNA localization patterns for genes encoding enzymes in the GA biosynthesis pathway (Fig. 1*F-K*). We observed that mRNAs encoding six enzymes leading to the synthesis of bioactive GA (*ent*-kaurene synthase A, *ent*-kaurene synthase B, *ent*-kaurene oxidase, *ent*-kaurenoic acid hydroxylase, GA 20-oxidase, and GA 3-oxidase) accumulated primarily in the giant suspensor region (Fig. 1*F-K*). This extends previous studies that showed that SRB suspensors are a rich source of GA (16), synthesize GA in cell-free extracts (17, 18), and contain GA 3-oxidase mRNA (6, 19).

We confirmed our SRB GA mRNA localization studies and expanded them to the closely related Common Bean by using (i) laser-capture micro-dissection (LCM) technology to collect SRB and Common Bean globular-stage embryo proper and suspensor regions, (ii) RNA-Seq for transcriptome profiling, and (iii) the Common Bean as a reference genome (20) (GEO accession GSE57537) (Fig. 1*L-Q*). The genome browser view illustrates that the up-regulation of GA biosynthesis enzyme mRNAs in the giant suspensor relative to the embryo proper was conserved in both of these bean species. By contrast, GA 2-oxidase mRNA, accumulated to very low, or non-detectable, levels in SRB and Common Bean globular-stage embryos (Table S1). Our data indicate that the synthesis of bioactive GA within giant bean suspensors (16) is primarily due to the spatially restricted accumulation of GA biosynthesis enzyme mRNAs.

We examined the temporal mRNA accumulation pattern of *GA 20-oxidase* during early SRB embryo development (Fig. 3*A-E*). GA 20-oxidase mRNA was first detected in the basal cell of a two-cell embryo shortly after fertilization, and then accumulated to high levels in the suspensor from pre-globular stage to heart stage (Fig. 3*B-E*), similar to what we observed for G564 mRNA (12). Later, GA 20-oxidase mRNA accumulated within the epidermis of the heart stage embryo proper (Fig. 3*E*). The GA 3-oxidase mRNA temporal accumulation pattern in SRB embryos (Fig. 4) was indistinguishable from that of GA 20-oxidase mRNA (Fig. 3*B-E*), as well as G564 mRNA (12). These results suggest that genes encoding GA biosynthesis enzymes are regulated by the same *cis*-regulatory elements as *G564*, and form part of a SRB suspensor gene regulatory network that is activated shortly after fertilization. To test this hypothesis, we used transgenic tobacco embryos to search for *cis*-regulatory elements required for *GA 20-oxidase* transcription within the SRB suspensor, similar to the approach that we used for *G564* (11, 12, 14).

**Fig. 3.**
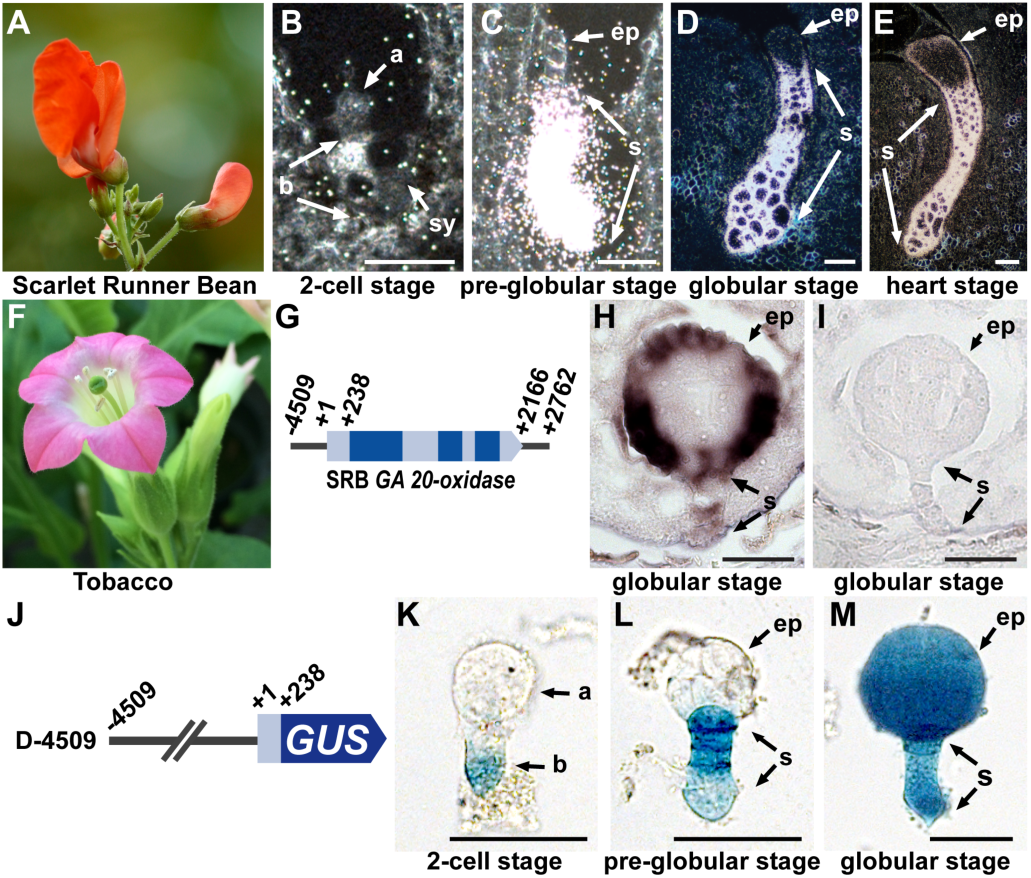
GA 20-oxidase mRNA localization in SRB embryos and *GA 20-oxidase* transcriptional activity during tobacco embryogenesis. (*A*) SRB flower (image taken from Fig. 1*A*). (*B-E*) Localization of GA 20-oxidase mRNA in developing SRB embryos: (*B*) two-cell stage, (*C*) pre-globular stage, and (*D*) globular stage (image taken from Fig. 1*J*), and (*E*) heart stage. (*F*) Tobacco flower. (*G*) Conceptual representation of the SRB *GA 20-oxidase* transgene introduced into tobacco. Dark blue boxes represent exons. Light blue boxes represent UTRs and introns. Numbers indicate positions relative to the transcription start site (+1). (*H-I*) Hybridization of SRB *GA 20-oxidase* anti-sense (*H*) and sense (*I*) probes to globular-stage transgenic tobacco embryos. (*J*) Conceptual representation of the *GA 20-oxidase/GUS* transgene introduced into tobacco. (*K-M*) GUS activity in transgenic tobacco embryos: two-cell stage (*K*), pre-globular stage (*L*), globular stage (*M*). Photographs were taken after 24-h GUS incubation for two-cell and pre-globular stage, and 1-h for globular stage. a, apical cell; b, basal cell; ep, embryo proper; s, suspensor; sy, synergid. (Scale bar: 50 µm.)

**Fig. 4.**
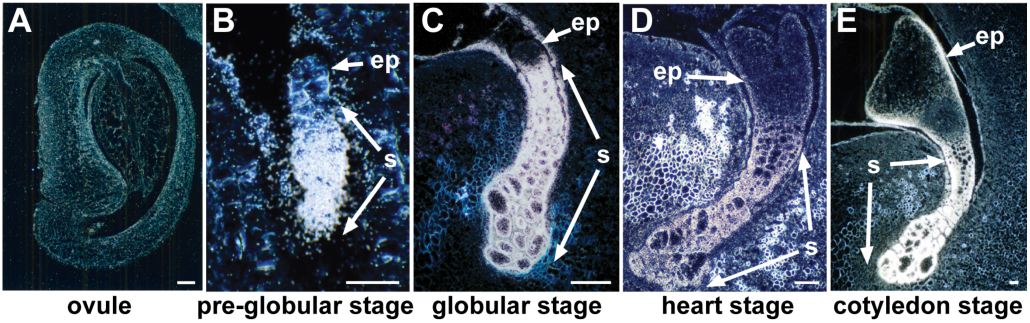
Localization of GA 3-oxidase mRNA in SRB ovule and embryos. (*A*) Ovule. (*B*) Pre-globular stage. (*C*) Globular stage (image taken from Fig. 1*K*). (*D*) Heart stage. (*E*) Cotyledon stage. (Scale bar: 50 µm.)

### SRB GA 20-oxidase mRNA accumulates within the suspensor of transgenic tobacco embryos

We transformed tobacco (Fig. 3*F*) with a 7.271-kb SRB *GA 20-oxidase* genomic fragment (Fig. 3*G*), and localized GA 20-oxidase mRNA in transgenic globular-stage embryos using *in situ* hybridization (Fig. 3*H* and *I*). The *GA 20-oxidase* 5’ and 3’ regions were 4,509 and 596 bp in length, respectively, and did not contain similarity to any known genes. SRB GA 20-oxidase mRNA localized within tobacco suspensor and embryo proper protodermal cells, the precursors to heart-stage epidermal cells (Fig. 3*H* and *I*), similar to the GA 20-oxidase mRNA accumulation pattern in SRB embryos (Fig. 3*E*). These results indicate that the GA 20-oxidase mRNA accumulation pattern is conserved during early embryo development in both tobacco and SRB.

### *GA 20-oxidase* expression within the suspensor is under transcriptional control

We introduced a chimeric SRB *GA 20-oxidase/β-glucuronidase (GUS)* gene into tobacco (Fig. 3*J*) and localized GUS enzyme activity in transgenic embryos to study *GA 20-oxidase* transcriptional regulation (Fig. 3*K-M*). The *GA 20-oxidase* region −4,509/+238 fused to *GUS* (D-4509) (Fig. 3*J*) first programmed GUS enzyme activity within the basal region of the two-cell tobacco embryo, followed by the entire suspensor at the pre-globular stage, and then to the globular-stage embryo proper (Fig. 3*K-M*). The GUS activity pattern was congruent with the localization of GA 20-oxidase mRNA in tobacco embryos driven by the entire *GA 20-oxidase* gene (Fig. 3*H* and *I*), as well as during SRB embryo development (Fig. 3*B-E*). These results indicate that (i) the temporal and spatial expression pattern of *GA 20-oxidase* is controlled primarily at the transcriptional level by sequences within the −4,509/+238 region (Fig. 3*J*) and (ii) the regulatory apparatus that controls *GA 20-oxidase* gene activity during early embryo development is conserved between SRB and tobacco.

### The *GA 20-oxidase* upstream region contains separate embryo proper and suspensor *cis*-regulatory regions

We generated 5’ deletions of the *GA 20-oxidase* −4,509 to +238 region, and analyzed GUS activity in transgenic tobacco embryos to identify sequences required for transcription in the globular embryo (Fig. 5). Progressively deleting sequences from −4,509 to −275 first caused a loss of GUS activity in the embryo proper, followed by loss of GUS activity in the suspensor (Fig. 5), indicating that there are separate embryo proper (−2,000 to −1,500) and suspensor (−450 to +238) *cis*-regulatory regions. The −450 to +238 *GA 20-oxidase* region appears to contain all the sequences required for transcription within the suspensor. Deletion to −275 reduced suspensor GUS activity to a barely detectable level, similar to that observed in the *GUS*-only negative control, which may be due to a low level of cryptic transcripts initiating in the vector sequence and reading through the reporter gene, as has been demonstrated for other plasmids (21). We conclude that sequences in the 175 bp region −450 to −275 are required for suspensor transcription, although additional downstream sequences to +238 might also be required.

**Fig. 5.**
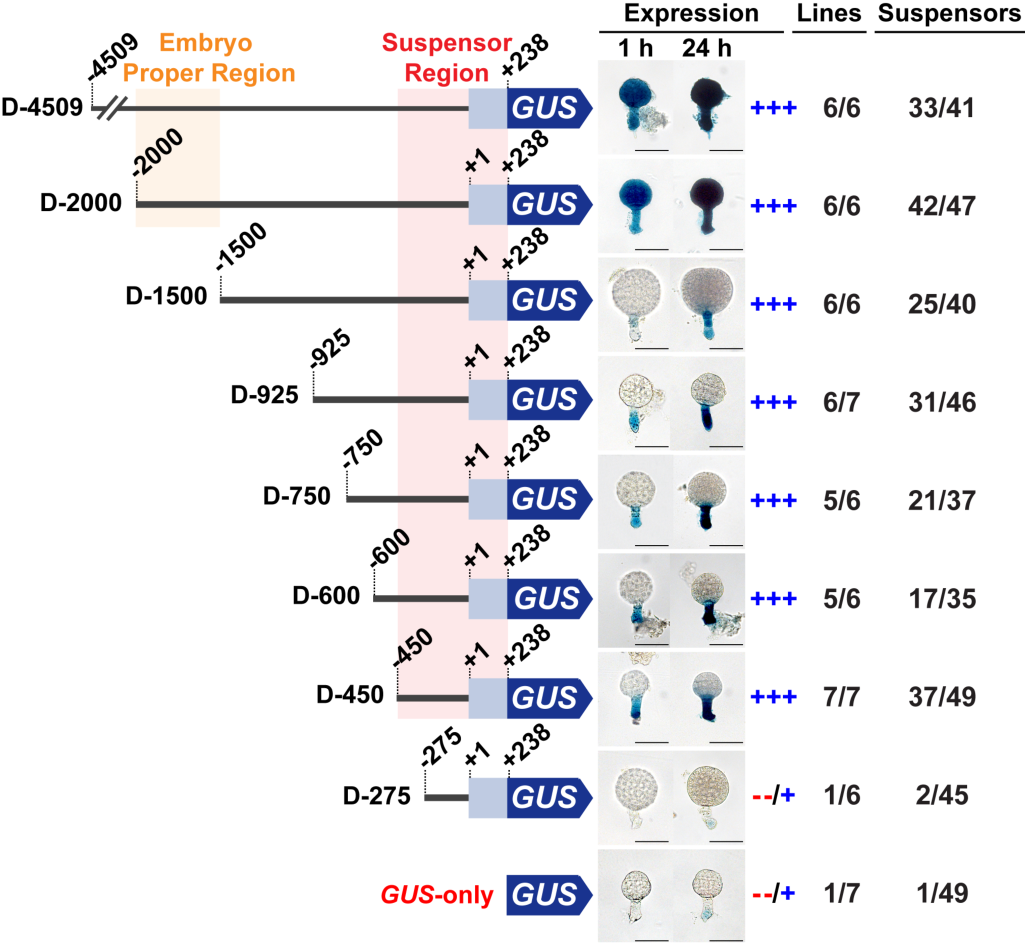
GUS activity in transgenic tobacco embryos containing 5’ deletions of the *GA 20-oxidase* upstream region. Conceptual representations of the constructs are to the left of each embryo. Dark blue arrows represent the *GUS* gene. Light blue boxes represent the *GA 20-oxidase* 5’ UTR. Numbers indicate positions relative to the *GA 20-oxidase* transcription start site (+1). Embryo proper and suspensor *cis*-regulatory regions are highlighted in orange and red, respectively. Expression levels were categorized as previously described (11, 14). +++ in the Expression column indicates that suspensor GUS activity was strong; that is, the majority of the suspensors with GUS activity at 24-h were GUS-positive at 2-h. −/+ in the Expression column indicates that suspensor GUS activity was weak; that is, the majority of the suspensors with GUS activity at 24-h were GUS-negative at 2-h. Numbers in the Lines column indicate the number of individual transformants displaying suspensor GUS activity over the total number of individual transformants analyzed. Numbers in the Suspensors column indicate the number of embryos displaying suspensor GUS activity by 24-h incubation over the total number of embryos analyzed. Photographs were taken after 1-h and 24-h GUS incubation. (Scale bar: 50 µm.)

### G564 suspensor *cis*-control motifs are present in the *GA 20 oxidase* −450 to +238 region

Because *GA 20-oxidase* has the same suspensor-specific expression pattern as *G564* (Fig. 1*E* and *J*) (12), we searched the *GA 20-oxidase* −450 to +238 region for the presence of known suspensor *cis*-regulatory elements that activate *G564* transcription: (i) the 10-bp motif (5’-GAAAAGCGAA-3’ with up to three non-adjacent mismatches), (ii) the Region 2 motif (5’-TTG(A/G)(A/G/T)AAT-3’), and (iii) the Fifth motif (5’-(A/G)AGTTA-3’) (14). Within the −450 to +238 region we identified eight predicted 10-bp motifs, three predicted Fifth motifs, and six Region 2 motifs allowing for one mismatch – except that the third nucleotide was not an A, as this nucleotide inactivates the Region 2 motif (11) (Fig. 6*A*).

**Fig. 6.**
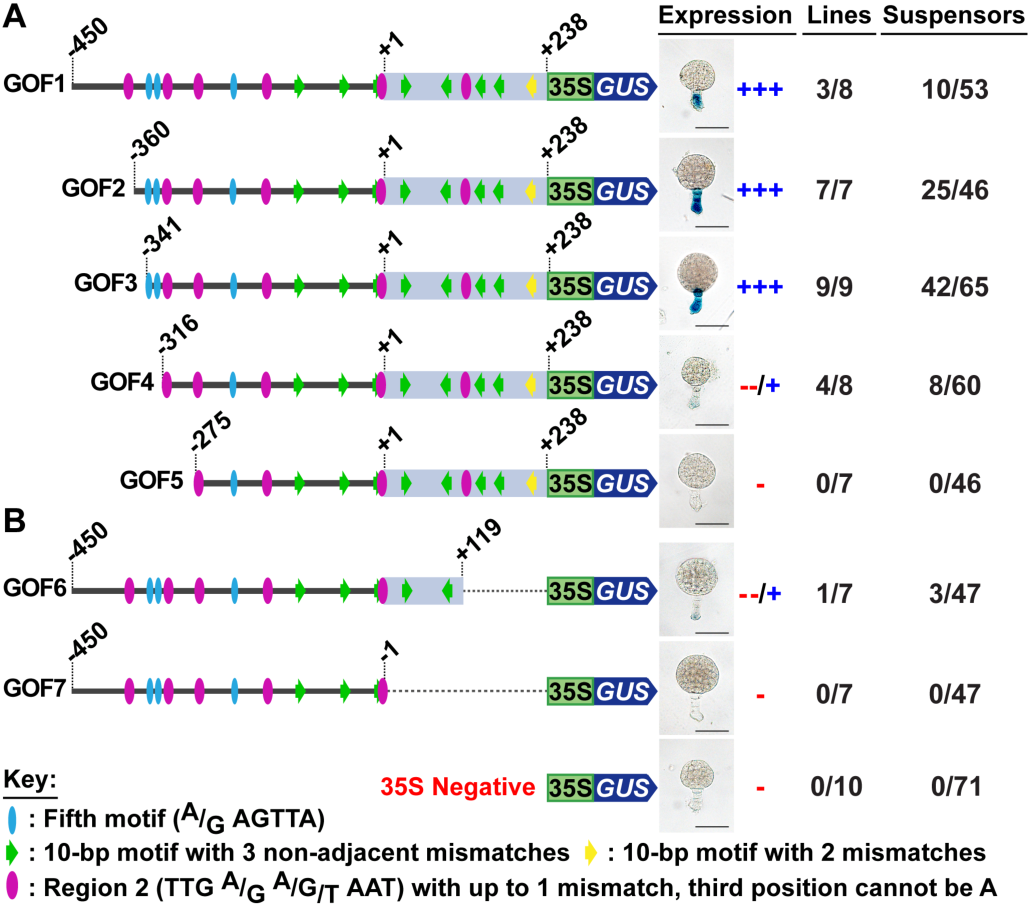
GUS activity in transgenic tobacco embryos containing 5’ (*A*) and 3’ (*B*) deletions of the *GA 20-oxidase* upstream region fused to the *CaMV 35S* minimal promoter/*GUS* gene. Conceptual representations of the constructs are to the left of each embryo. Yellow and green arrows, purple ovals, and blue ovals represent the 10-bp motif, Region 2 motif, and Fifth motif, respectively, with their sequences defined in the key. Dark blue arrows represent the *GUS* gene. Light blue boxes represent the *GA 20-oxidase* 5’ UTR. Green boxes represent the *CaMV 35S* minimal promoter. Numbers indicate positions relative to the *GA 20-oxidase* transcription start site (+1). Expression levels were categorized previously described (11, 14). +++ in the Expression column indicates that suspensor GUS activity was strong; that is, the majority of the suspensors with GUS activity at 24-h were GUS-positive at 2-h. −/+ in the Expression column indicates that suspensor GUS activity was weak; that is, the majority of the suspensors with GUS activity at 24-h were GUS-negative at 2-h. Minus (-) in the Expression column indicates no detectable suspensor GUS activity at 24-h. Numbers in the Lines column indicate the number of individual transformants displaying suspensor GUS activity over the total number of individual transformants analyzed. Numbers in the Suspensors column indicate the number of embryos displaying suspensor GUS activity by 24-h incubation over the total number of embryos analyzed. Photographs were taken after 24-h GUS incubation. (Scale bar: 50 µm.)

### The *GA 20-oxidase* upstream region −341 to −316 is required for suspensor transcription

To determine which of the predicted *G564* motifs might be required for *GA 20-oxidase* suspensor transcription, we performed additional 5’ deletions within a gain-of-function (GOF) construct containing the *GA 20-oxidase* −450 to +238 upstream region fused to a *Cauliflower Mosaic Virus (CaMV) 35S* minimal promoter and *GUS* (GOF1) (Fig. 6*A*). This construct programmed high levels of GUS activity specifically within the suspensor (Fig. 6*A*) as predicted from our initial 5’ deletion analysis (Fig. 5), confirming that all of the sequences required for *GA 20-oxidase* suspensor transcription are present within the −450 to +238 region. Deletions to −360 (GOF2) and −341 (GOF3) did not affect GUS activity, whereas deletion to −316 (GOF4) decreased GUS activity significantly within the suspensor (Fig. 6*A*). Together, these data show that the region −341 to −316 is required for full transcriptional activity in the suspensor and contains two predicted Fifth motifs, which may be functional.

### The *GA 20-oxidase* upstream region +119 to +238 is required for suspensor transcription

To determine whether sequences downstream of −341 to −316 are also required for *GA 20-oxidase* suspensor transcription, we performed 3’ deletions of the −450 to +238 GOF1 construct (Fig. 6*B*). A 3’ deletion to +119, or half of the 5’ UTR, (GOF6), resulted in a significant decrease in GUS activity (Fig. 6*B*), indicating that sequences in the +119 to +238 *GA 20-oxidase* region were required for suspensor transcription. Further 3’ deletion (GOF7), did not affect significantly *GA 20-oxidase* suspensor transcription (Fig. 6*B*). We conclude that one or more suspensor *cis*-regulatory elements are present within the +119 to +238 region, which includes three predicted 10-bp motifs and one predicted Region 2 motif (compare GOF5 and GOF6 in Fig. 6).

### 10-bp and Region 2 motifs are required for *GA 20-oxidase* suspensor transcription

We carried out site-directed mutagenesis and 3’ deletion experiments within the GOF2 construct to determine which of the predicted 10-bp and Region 2 motifs in the +119 to +238 region were functional (Fig. 7). Mutation of the predicted Region 2 motif in this region (M1) caused a significant decrease in suspensor transcription, similar to the level of GUS activity observed when the +119 to +238 region containing this motif was deleted (GOF6) (Fig. 7). This result indicates that the Region 2 motif is required for *GA 20-oxidase* suspensor transcription.

**Fig. 7.**
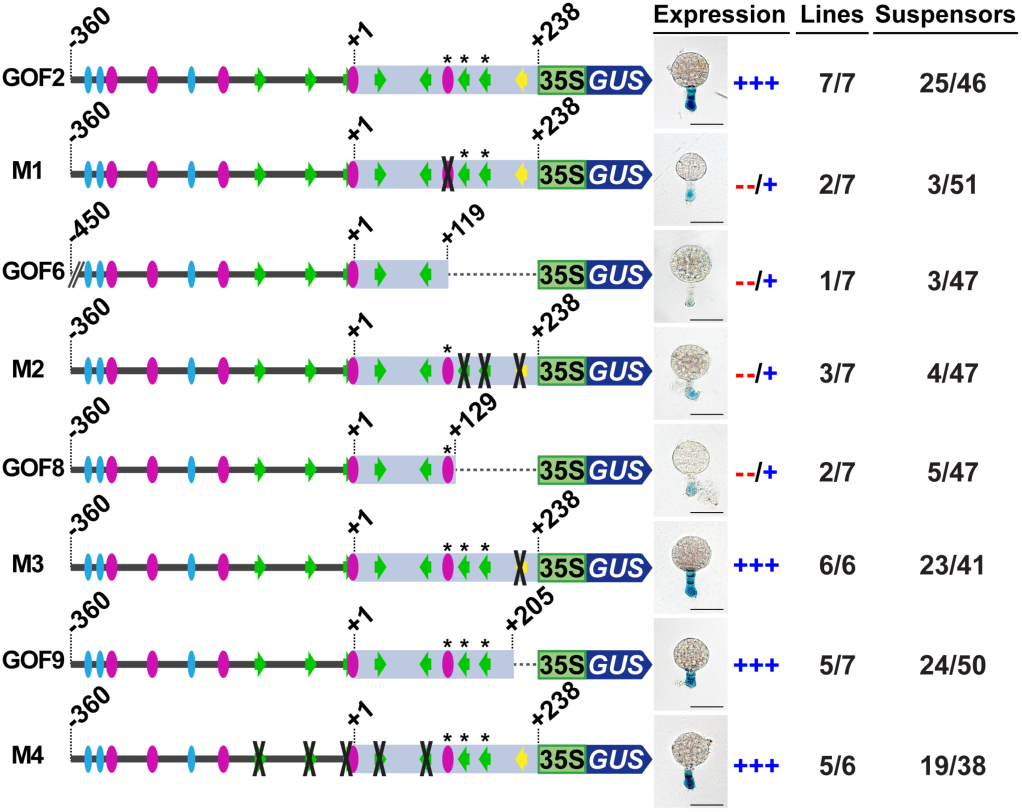
GUS activity in transgenic tobacco embryos containing site-directed mutations or 3’ deletions within the *GA 20-oxidase* +119 to +238 region. Conceptual representations of the constructs are to the left of each embryo. Yellow and green arrows, purple ovals, and blue ovals represent the 10-bp motif, Region 2 motif, and Fifth motif, respectively. Black crosses indicate mutations of the indicated motifs. Asterisks indicate the motifs found to be required for suspensor transcription. Dark blue arrows represent the *GUS* gene. Light blue boxes represent the *GA 20-oxidase* 5’ UTR. Green boxes represent the *CaMV 35S* minimal promoter. Numbers indicate positions relative to the *GA 20-oxidase* transcription start site (+1). Expression levels were categorized as described previously (11, 14). +++ in the Expression column indicates that suspensor GUS activity was strong; that is, the majority of the suspensors with GUS activity at 24-h were GUS-positive at 2-h. −/+ in the Expression column indicates that suspensor GUS activity was weak; that is, the majority of the suspensors with GUS activity at 24-h were GUS-negative at 2-h. Minus (-) in the Expression column indicates no detectable suspensor GUS activity at 24-h. Numbers in the Lines column indicate the number of individual transformants displaying suspensor GUS activity over the total number of individual transformants analyzed. Numbers in the Suspensors column indicate the number of embryos displaying suspensor GUS activity by 24-h incubation over the total number of embryos analyzed. Photographs were taken after 24-h GUS incubation. The GOF2 and GOF6 constructs are reproduced from Fig. 6 for comparative purposes. (Scale bar: 50 µm.)

To determine whether the three predicted 10-bp motifs within this region were also required, we mutated and deleted these motifs (M2 and GOF8, Fig. 7). Both mutation (M2) and deletion (GOF8) of the three predicted 10-bp motifs decreased GUS activity significantly (Fig. 7), indicating that at least one, or more, of the three 10-bp motifs were required for suspensor transcription (Fig. 7). We hypothesized that the proximal 10-bp motif (Fig. 7, yellow arrow) was required because it most closely resembled the consensus *G564* 10-bp motif, having only two mismatches instead of three. However, neither mutation (M3) nor deletion (GOF9) of this 10-bp motif affected GUS activity (Fig. 7). We conclude that at least one, or both, of the remaining 10-bp motifs in the +129 to +205 region are required for suspensor transcription.

Because the *G564* suspensor *cis*-regulatory module requires three copies of the 10-bp motif (14), we asked whether any predicted 10-bp motifs in the region upstream of +129 were also required for *GA 20-oxidase* suspensor transcription. Mutating the remaining five predicted 10-bp motifs (M4) had no effect on suspensor GUS activity (Fig. 7), indicating that probably both 10-bp motifs in the +129 to +205 UTR region are required for *GA 20-oxidase* suspensor transcription in addition to the Region 2 motif.

### The Fifth motif and an additional Region 2 motif are required for *GA 20-oxidase* suspensor transcription

Two Fifth motifs were predicted in the −341 to −316 *GA 20-oxidase* upstream region, which was also required for suspensor transcription (compare GOF3 and GOF4, Figs. 6*A* and 8*A*). Site-directed mutagenesis was performed within the GOF2 construct to determine whether these predicted Fifth motifs at −336 to −341 and −329 to −324 were functional (Fig. 8*B*). Mutation of both predicted Fifth motifs in this region by either transversional mutagenesis (M5) or adenine substitution (M6) did not affect suspensor GUS activity (Fig. 8*B*). Because (i) the −341 to −316 region was required for suspensor transcription (compare GOF3 and GOF4, Figs. 6*A* and 8*A*), and (ii) a Fifth motif is essential for *G564* suspensor transcription (14), we searched the −341 to −316 region again for a Fifth motif allowing for one mismatch at any position in the consensus sequence (Fig. 2). We identified an additional Fifth motif (asterisk in GOF2 at −317 to −312) that overlapped a predicted Region 2 motif in the opposite orientation (Fig. 8*B*). Previously, we showed that *G564* motif orientation did not affect suspensor transcription (11).

**Fig. 8.**
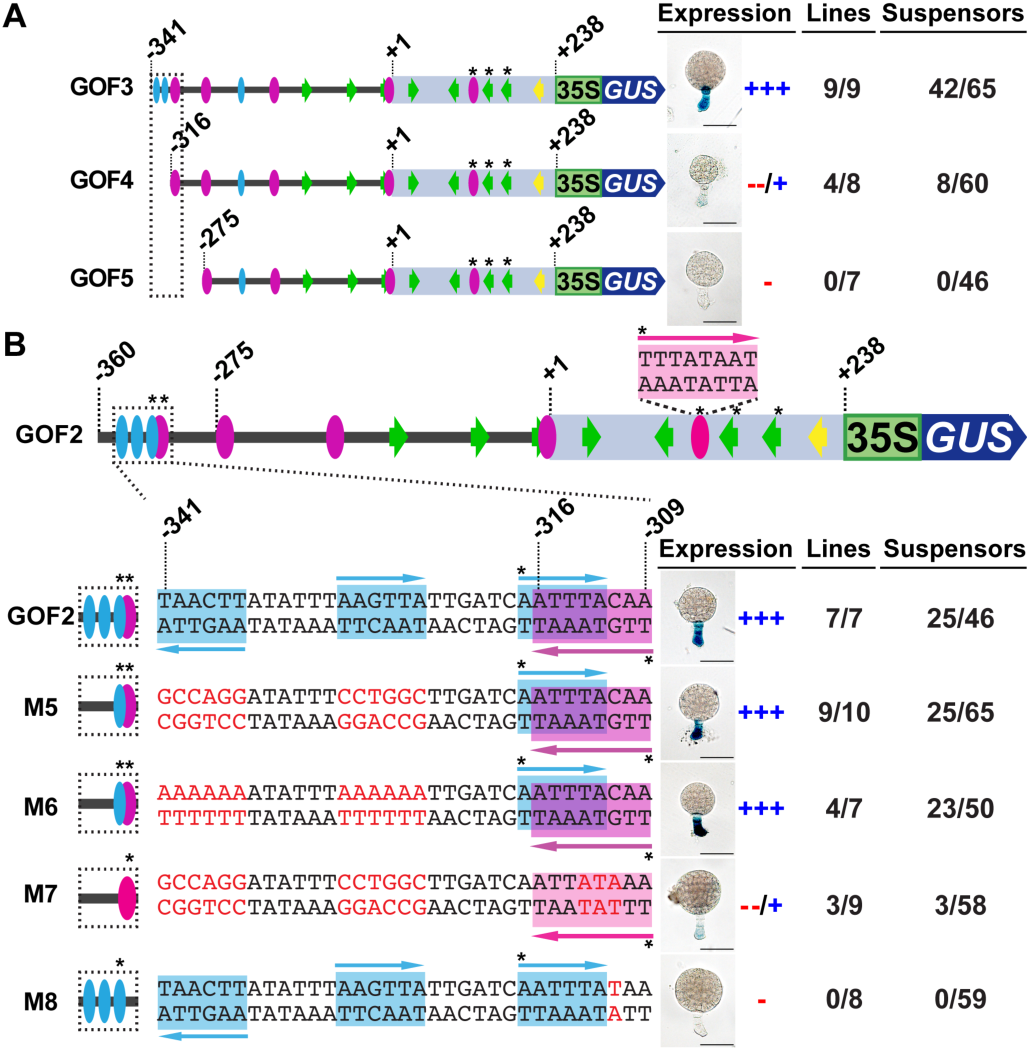
GUS activity in transgenic tobacco embryos containing 5’ deletions (*A*) and site-directed mutations (*B*) within the *GA 20-oxidase* −341 to −309 region. (*A*) 5’ GOF deletion constructs reproduced from Fig. 6*A* for comparative purposes. (*B*) Mutated constructs focusing on the −341 to −309 region. The sequence of the −341 to −309 region of each construct is shown to the left of each embryo. A conceptual representation of the predicted motifs in this region is shown in the dashed box to the left of the sequence. Region 2 and Fifth motif sequences are highlighted in purple and blue, respectively. The Region 2 motif sequence located in the 5’ UTR is shown above the GOF2 construct in pink. Arrows indicate the motif sequence orientation. Mutation sequences are shown in red font. The Fifth motif at −317 to −312 has one mismatch relative to the *G564* consensus sequence (14). Yellow and green arrows, purple ovals, and blue ovals represent the 10-bp motif, Region 2 motif, and Fifth motif, respectively. Asterisks indicate the motifs found to be required for suspensor transcription. Dark blue arrows represent the *GUS* gene. Light blue boxes represent the *GA 20-oxidase* 5’ UTR. Green boxes represent the *CaMV 35S* minimal promoter. Numbers indicate positions relative to the *GA 20-oxidase* transcription start site (+1). Expression levels were categorized as described previously (11, 14). +++ in the Expression column indicates that suspensor GUS activity was strong; that is, the majority of the suspensors with GUS activity at 24-h were GUS-positive at 2-h. −/+ in the Expression column indicates that suspensor GUS activity was weak; that is, the majority of the suspensors with GUS activity at 24-h were GUS-negative at 2-h. Minus (-) in the Expression column indicates no detectable suspensor GUS activity at 24-h. Numbers in the Lines column indicate the number of individual transformants displaying suspensor GUS activity over the total number of individual transformants analyzed. Numbers in the Suspensors column indicate the number of embryos displaying suspensor GUS activity by 24-h incubation over the total number of embryos analyzed. GOF2 deletion construct is reproduced from Fig. 6*A* for comparative purposes. Photographs were taken after 24-h GUS incubation. (Scale bar: 50 µm.)

To determine whether the Fifth motif at −317 to −312 was required for *GA 20-oxidase* suspensor transcription, site-directed mutagenesis was performed on the GOF2 construct (M7) (Fig. 8*B*). In order to mutate the Fifth motif and leave the overlapping Region 2 motif intact, we had to substitute the predicted Region 2 motif at −309 to −316 (5’-TTGTAAAT-3’) with the functional Region 2 motif from the 5’ UTR (5’-TTTATAAT-3’), which had a slightly different sequence (M7, Fig. 8*B*). This caused a mutation (red nucleotides) in the overlapping predicted Fifth motif while keeping the Region 2 motif intact (M7, Fig. 8*B*). Mutation of all three predicted Fifth motifs in the −341 to −316 region (M7) resulted in a significant decrease in suspensor transcription (M7, Fig. 8*B*), consistent with the results of GOF4 that deleted the −341 to −316 region, including the first nucleotide of the −317 to −312 Fifth motif (Figs. 6*A* and 8*A*). Because mutating the two predicted Fifth motifs at −336 to −341 and −329 to −324 had no effect on suspensor GUS activity (M5 and M6, Fig. 8*B*), the Fifth motif at −317 to −312 is required for suspensor transcription.

To determine whether the overlapping predicted Region 2 motif at −309 to −316 was also required for *GA 20-oxidase* suspensor transcription, we mutated this motif within the GOF2 construct by changing the third nucleotide (−311) from G to A, leaving the Fifth motif intact (M8, Fig. 8*B*). Previously, we demonstrated that this mutation renders the Region 2 motif nonfunctional in the *G564* suspensor *cis*-regulatory module (11). Similar to *G564*, mutating a single nucleotide in the Region 2 motif at −309 to −316 (M8) abolished suspensor GUS activity (Fig. 8*B*). Thus, the Region 2 motif at −309 to −316 is required for suspensor transcription, in addition to the Region 2 motif within the 5’ UTR. Together, these results indicate that both the Fifth motif and the Region 2 motif in the −317 to −309 *GA 20-oxidase* upstream region are essential for transcription within the suspensor, and explain the loss of suspensor GUS activity using the −275 deletion (D-275) and gain-of-function (GOF5) constructs (Figs. 5, 6*A*, and 8*A*).

### Functional suspensor motifs are conserved in the Common Bean *GA 20-oxidase* gene

We examined the Common Bean *GA 20-oxidase* gene region for suspensor motifs because it is nearly identical to SRB *GA 20-oxidase* in (i) structure (Fig. S1*A*), (ii) sequence (Fig. S1*B*), and (iii) expression pattern (Fig. 1*P*). We found sequences identical to the functional SRB *GA 20-oxidase* suspensor motifs in the Common Bean *GA 20-oxidase* gene region (Fig. S1). The sequence, order, orientation, and spacing (with the exception of small indels) of the functional motifs were identical in these bean species, suggesting that the Common Bean *GA 20-oxidase* gene also utilizes the 10-bp, Region 2, and Fifth motifs for transcription within its giant suspensor.

## Discussion

The giant SRB suspensor (Fig. 1) has been used for over four decades as a model system for investigating the physiological and cellular events that occur in this unique embryonic region, and what role it plays in early embryo development (5, 10, 22, 23). We have been using SRB suspensors to dissect the regulatory processes required for the region-specific transcription of genes within the embryo shortly after fertilization (6, 11, 12, 14). We identified a large number of genes, including *G564*, which are expressed specifically within the SRB suspensor using a variety of genomic approaches (6, 12). Genes encoding all major enzymes of the GA biosynthetic pathway (*ent-kaurene synthase A*, *ent-kaurene synthase B*, *ent-kaurene oxidase*, *ent-kaurenoic acid hydroxylase*, *GA 20-oxidase* and *GA 3-oxidase*) are also expressed specifically within giant SRB suspensors (Fig. 1) and, at least two (*GA 20-oxidase* and *GA 3-oxidase*), and probably all, are activated in the basal region of the post-fertilized embryo similar to *G564* (Fig. 3*B, C* and Fig. 4*B*) (12). Our results suggest that genes encoding GA biosynthesis enzymes, and others, such as *G564*, are organized into a genetic regulatory network (9) that (i) is activated after the division of the SRB embryo into the apical and basal regions and (ii) functions exclusively within the suspensor. This model predicts that genes operating within the suspensor genetic regulatory network share a common *cis*-regulatory module that is responsible for activating genes within the suspensor. The results presented in this paper support this model, showing that both *G564* and *GA 20-oxidase* genes require the same *cis*-regulatory motifs in order to be activated within the suspensor region (Fig. 9).

**Fig. 9.**
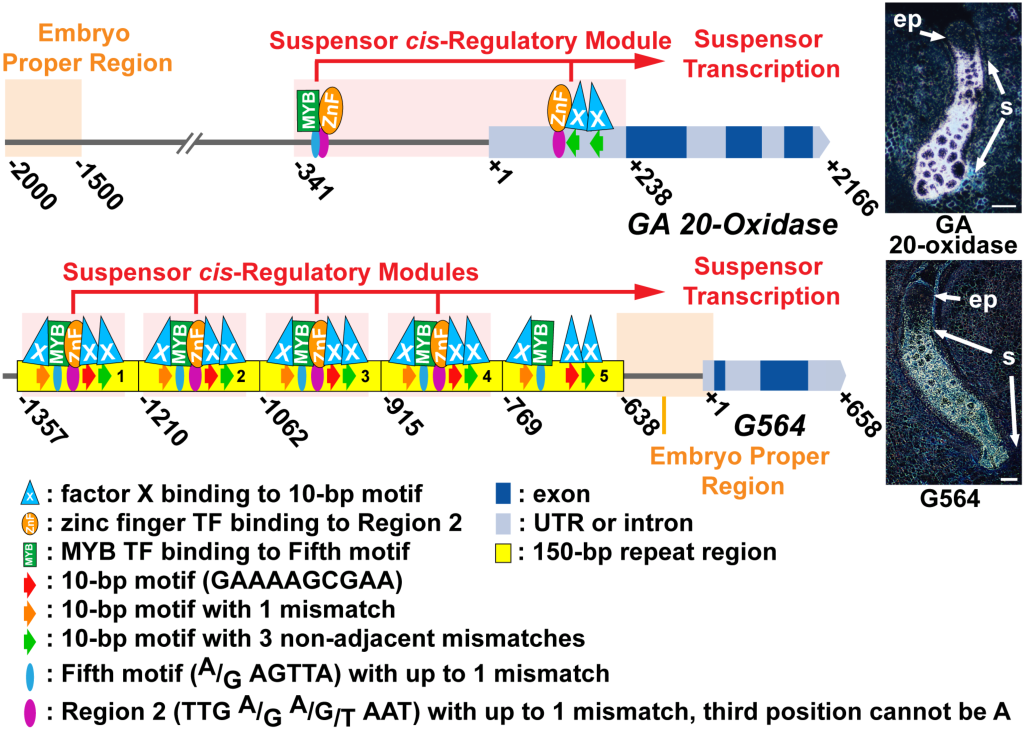
Models of SRB *GA 20-oxidase* and *G564* suspensor *cis*-regulatory module organization. Embryo proper and suspensor *cis*-regulatory regions are highlighted in orange and red, respectively. MYB and zinc finger (ZnF) are candidate transcription factors that bind to the Fifth and Region 2 motifs, respectively. GA 20-oxidase mRNA *in situ* image was reproduced from Fig 1*J*. G564 mRNA *in situ* image was reproduced from Weterings *et al*. (12). Data used to generate these models were taken from previously published papers from our lab (11, 14) and the results presented here.

### The SRB *GA 20-oxidase* upstream region is organized into different modules that direct expression to the suspensor and embryo proper

*GA 20-oxidase* contains two discrete regions that activate transcription in the suspensor and embryo proper during embryogenesis (Figs. 5 and 9). One region (−341 to +238) activates transcription within the suspensor following fertilization (Fig. 6*A*). The second region (−2000 to −1500) activates transcription later in embryo development within embryo proper epidermal cells (Fig. 5). This bi-modular organization of suspensor and embryo proper control regions is similar to that of *G564*, except that the spatial distribution of these modules differs (Fig. 9) (11, 12).

The *GA 20-oxidase* suspensor module activates transcription uniformly over cells of the entire suspensor, in contrast with the cell-specific embryo proper module (Fig. 5). Our deletion and mutagenesis experiments with the −341 to +238 suspensor control region (Figs. 5-8) did not uncover sub-regions required for transcription within specific suspensor cell types, such as the (i) hypophysis adjacent to the embryo proper that is derived from the embryo apical cell and (ii) enlarged basal cell that forms connections with the seed coat and is derived from the embryo basal cell. This suggests that the *GA 20-oxidase* gene is activated within suspensor cells using the same regulatory processes irrespective of position or cell lineage.

The *GA 20-oxidase* suspensor control region activates transcription within the basal cell of the two-cell embryo (Fig. 3*B* and *K*), and in all suspensor cells by the late globular stage of development (Figs. 3*B-D, K-M*). We did not observe any shift in this temporal sequence of transcriptional events in our deletion and mutagenesis experiments. Nor did we observe any ectopic activation of transcription within the globular-stage embryo proper region by deleting, or mutating, parts of the suspensor control module (Figs. 5-8). These results suggest the absence of both temporal and negative *cis*-regulatory elements within the suspensor regulatory region. Thus, the regulation of *GA 20-oxidase* within the SRB suspensor is controlled by positive elements that activate transcription shortly after fertilization, and within daughter suspensor cells as they form during embryogenesis.

The organization of regulatory sequences within the *GA 20-oxidase* suspensor control region is relatively simple compared with those that activate storage protein gene transcription within the mature embryo proper. Storage protein gene embryo-proper control regions contain (i) territory-specific modules regulating transcription within embryo-proper sub-regions, such as the axis and cotyledons [e.g. *Kti3* (2), *β-Phaseolin* (24)], (ii) temporal *cis*-regulatory elements [e.g. *β-Phaseolin* (24), *β-conglycinin* (25)], and (iii) repressor elements [e.g. *Glycinin* (26), *β-Phaseolin* (24)]. This difference in regulatory architecture most likely results from the fact that the suspensor is a highly specialized, terminally differentiated embryonic region with few distinct cell types and degenerates later in development, in contrast with the more-complex embryo proper that contains many functionally distinct embryonic territories that give rise to the mature plant following seed germination.

### At least five *cis*-regulatory elements are required to activate *GA 20-oxidase* suspensor transcription

Within the SRB *GA 20-oxidase* −341 to +238 suspensor control region, we identified 17 short sequences similar to the motifs that are required to activate *G564* in the suspensor – the 10-bp motif, Region 2 motif, and the Fifth motif (Figs. 2 and 6) (11, 14). Deletion and site-directed mutagenesis experiments demonstrated only five of these sequences are functional and required for *GA 20-oxidase* suspensor transcription – two Region 2 motifs, a Fifth motif, and two 10-bp motifs – demonstrating that both *G564* and *GA 20-oxidase* utilize the same suspensor *cis*-control elements in support of our original hypothesis (Figs. 7 and 8). All five of these motifs are conserved at similar positions within the Common Bean *GA 20-oxidase* gene region (Fig. S1). The remaining candidate motifs are not required for suspensor transcription (Figs. 7 and 8), illustrating the need to functionally dissect *cis*-regulatory modules to understand how they operate rather than relying solely on motif similarity from computer predictions.

The Fifth motif and Region 2 motif in the *GA 20-oxidase* −317 to −309 upstream region overlap (Figs. 8*B* and 9), and mutating either motif, without disrupting the other, leads to a decrease or complete loss of suspensor transcriptional activity (Fig. 8*B*). This unique regulatory element organization is not without precedent as functional overlapping motifs have been identified in several animal enhancer regions (27-29). By contrast, two 10-bp motifs and a second Region 2 motif are located in the *GA 20-oxidase* 5’ UTR and do not overlap (Fig. 9). Other plant genes contain positive transcriptional control elements in their 5’ UTRs (30, 31). Together, these data indicate that transcription factors must bind to motifs within the suspensor regulatory module that reside both upstream and downstream of the transcription start site in order to activate *GA 20-oxidase* transcription in the suspensor.

### Suspensor-specific gene transcription is generated by a flexible arrangement of *cis*-regulatory motifs

The results presented here on the organization of the *GA 20-oxidase* suspensor *cis*-regulatory module, and those carried out previously with *G564* (11, 14), provide a unique opportunity to compare the architecture of two suspensor *cis*-regulatory modules. The *G564* suspensor module is composed of three 10-bp motifs, a Region 2 motif, and a Fifth motif all tightly clustered within a 47-bp DNA region with little spacing between motifs (Fig. 9) (14). The *G564* suspensor *cis*-regulatory module is repeated five times in the *G564* upstream region, and each repeat is able to function individually except for repeat five as it lacks an intact Region 2 motif (Fig. 9) (11, 14).

By contrast, the single *GA 20-oxidase* suspensor module is larger, 579 bp in length, composed of two 10-bp motifs, two Region 2 motifs, and a Fifth motif divided between upstream and 5’ UTR regions (Fig. 9). Thus, the number, order, and spacing of suspensor motifs differ between the *G564* and *GA 20-oxidase cis*-regulatory modules even though they lead to the same output – suspensor-specific transcription within the early plant embryo. This suggests that the suspensor *cis*-regulatory module most closely resembles a billboard-type model of control element organization in which motif positions can vary between genes that program transcription to the same developmental state, as compared with an enhanceosome-type model that requires fixed motif positions for similarly expressed genes (32).

### Candidate transcription factors have been identified that bind to specific motifs within the suspensor control module

The *G564* and *GA 20-oxidase* suspensor *cis*-regulatory module can activate transcription within the suspensor of divergent plant embryos – including SRB, Common Bean, tobacco, and *Arabidopsis* (Figs. 1 and 3) (5, 11, 14). This suggests that the suspensor *cis*-regulatory module operates within a highly conserved regulatory network that utilizes a set of transcription factors that is shared between the suspensors of these plant species.

What transcription factors bind to the 10-bp, Region 2, and Fifth motifs that are required for the suspensor control module to function (Fig. 9)? We previously showed that the *G564* Fifth motif resembles the canonical sequence of a MYB transcription factor binding site, and yeast one-hybrid experiments with *Arabidopsis* transcription factors showed that several MYB transcription factors bind to the *G564* suspensor control module to activate transcription within yeast cells (14, 33). Although we have yet to identify the specific MYB transcription factor that binds to the Fifth motif, there are a number of SRB MYB transcription factors that are encoded by suspensor-specific mRNAs (RNA-Seq dataset GEO accession GSE57537) that are ideal candidates.

We searched the *Arabidopsis* DNA affinity purification sequencing (DAP-Seq) database (34) and identified an *Arabidopsis* C2H2-type zinc finger transcription factor (AT2G41835) that binds to the sequence 5’-TTGA(A/G)AA-3’ which is nearly identical to the Region 2 sequence 5’-TTG(A/G)(A/G/T)AAT-3’ (Fig. 2). Our yeast one-hybrid screen with *Arabidopsis* transcription factors showed that zinc finger transcription factors could activate the *G564 cis*-regulatory module, although AT2G41835 was not represented in the library (14, 33). We searched the Common Bean genome database (20) and identified a gene (Phvul.007G253000) that (i) encodes a protein that closely resembles the *Arabidopsis* AT2G41835 C2H2-type zinc finger transcription factor (ii) has 99% identity to a SRB genomic sequence contig (g017197_00035), (iii) is represented by SRB suspensor ESTs (GenBank IDs GD428417.1 and GD420985.1), and (iv) is expressed in both the embryo proper and suspensor of Common Bean and SRB globular-stage embryos, although to an elevated level within the suspensor (RNA-Seq dataset GEO accession GSE57537). This suggests that the Phvul.007G253000 C2H2-type zinc finger transcription factor is an excellent candidate for interacting with the Region 2 motif (Fig. 9). Interestingly, a close relative of the Phvul.007G253000 C2H2-type zinc finger transcription factor is represented in maize egg cell and zygote mRNA populations (35), suggesting that it might be present prior to fertilization in SRB and Common Bean as well.

The transcription factor that interacts with the 10-bp motif is not represented in the DAP-Seq database, or other plant transcription factor databases, and remains unknown (Fig. 9).

Together, the data presented here and elsewhere (11, 12, 14) demonstrate that the SRB *G564* and *GA 20-oxidase* genes are activated transcriptionally within the suspensor by the same *cis*-regulatory module. This strongly suggests that *G564*, *GA 20-oxidase*, and, most likely, other genes in the GA biosynthesis pathway (Fig. 1) form part of a suspensor genetic regulatory network. The precise nature of this regulatory network, and how it is activated specifically within the embryo basal region shortly after fertilization remain to be determined.

## Materials and Methods

### Plant Materials

Plants of the day-neutral SRB cultivar “Hammond’s Dwarf Red Flower” (Vermont Bean Seed Company, Fair Haven, VT) were grown in a greenhouse as described previously (12). Open flowers were pollinated by hand using a watercolor brush. Hand-pollinated flowers were tagged, and seeds were harvested from 2-8 days after pollination (DAP), as described previously (12). Common Bean seeds (Accession G19833) were obtained from Dr. Phillip E. McClean at North Dakota State University. Common Bean plants were grown under the same conditions as SRB for one month and then moved to a growth chamber with an 8-hr-light/16-hr-dark cycle to induce flowering. Seeds 1.6 − 2.0 mm in length were collected at 5-6 DAP.

### Radioactive *In Situ* Hybridization Analysis

Radioactive *in situ* hybridization studies were performed as described previously (12). Briefly, SRB seeds (2-8 DAP) were harvested, and seeds were cut at their chalazal ends before fixing to enhance penetration of the fixative. SRB seeds were fixed overnight at 4°C in 1% glutaraldehyde, 0.1 M sodium phosphate buffer, pH 7.0, and 0.1% Triton X-100. Fixed seeds were dehydrated, cleared, and embedded in paraffin. Eight-µm sections were hybridized with ^33^P-labeled antisense RNA probes. Probes were generated from cDNA clones made from micro-dissected 6 DAP suspensor regions of globular-stage SRB embryos (12). These cDNA clones corresponded to Common Bean GA biosynthesis enzyme genes: *ent-kaurene synthase A* (Phvul.001G152100), *ent-kaurene synthase B* (Phvul.005G048500), *ent-kaurene oxidase* (Phvul.005G183600), *ent*-*kaurenoic acid hydroxylase* (Phvul.006G123500), *GA 20-oxidase* (Phvul.010G087500), and *GA 3-oxidase* (Phvul.009G097100). After hybridization and emulsion development, sections were stained with 0.05% toluidine blue in 0.05% borate solution. Photographs were taken using dark-field illumination with a compound microscope (Olympus BH2; Olympus Corp., Lake Success, NY). The photographs were digitized, adjusted for optimum silver grain resolution using the KPT-Equalizer program (Metacreations Corp., Carpinteria, CA), and assembled in Adobe Photoshop CS5.1 (San Jose, CA). Probe sequences are listed in Table S2.

### Non-Radioactive *In Situ* Hybridization Analysis

Non-radioactive *in situ* hybridization with transgenic tobacco embryos was carried out using digoxigenin-labeled riboprobes (36). Briefly, transgenic tobacco seeds 10 DAP were harvested and fixed overnight at 4°C in 10% formalin/5% acetic acid/50% ethanol (37). Fixed seeds were dehydrated, cleared, and embedded in paraffin using a Leica ASP300S Tissue Processor. Six-µm sections were hybridized to sense or antisense digoxigenin-labeled riboprobes overnight. Probes were generated from a *GA 20-oxidase* cDNA clone containing the region +182 to +2166, which was isolated from micro-dissected suspensor regions of 6 DAP globular-stage SRB embryos (12). Photographs were taken using bright-field illumination with a compound microscope (Leica 5000 B).

### Bright-Field Microscopy

SRB and Common Bean seeds were fixed overnight at 4°C in 1% glutaraldehyde, 0.1 M sodium phosphate buffer, pH 7.0, and 0.1% Triton X-100. Fixed seeds were dehydrated and cleared. SRB seeds were embedded in Spurr’s plastic resin (38) (Polysciences, Warrington, PA). One-µm sections were stained for 18-20 min at 42°C with 0.05% toluidine blue in 0.05% borate solution. Photographs were taken using bright-field illumination with a compound microscope (Leica 5000 B). Common bean seeds were embedded in paraffin. Six-µm sections were stained for 1.5 min at 42°C with 0.1% toluidine blue. Photographs were taken using bright-field illumination with a compound microscope (Leica 5000 B).

### Plant Transformation

Tobacco (*Nicotiana tabacum* cultivar SR1) plants were transformed and regenerated using the leaf disk procedure (39). Each individual transformant was checked for T-DNA insertion by PCR and/or sequencing analysis. At least six independent transformants were generated for each construct. A total of 27 different constructs and 193 individual tobacco transformants were generated in order to carry out this study.

### GUS Histochemical Assay

Transgenic tobacco seeds were harvested 8 days after pollination (DAP). Globular-stage embryos were hand-dissected from seeds and assayed for GUS activity after 1-h, 2-h and 24-h at 37°C as described previously (11). Embryos were photographed under bright-field illumination using a compound microscope (Leica 5000 B; Leica). T1 seeds from GUS-negative lines were tested for kanamycin-resistant segregation after selfing to confirm that the T-DNA was not silenced. In total, 1,330 individual globular-stage embryos were assayed for GUS activity in order to generate the results reported in this study.

### 5’ Deletion Constructs

A *GA 20-oxidase* genomic clone (GenBank accession no. FJ535441) was digested with EcoRI and HindIII, and a 5.5 kb fragment containing the upstream sequence and the first exon was cloned into EcoRI- and HindIII-digested pBlueScriptII (Stratagene) generating plasmid pBSII PCS336 EH 5.5kb. To isolate the *GA 20-oxidase* upstream region, PCR was performed using the pBSII PCS336 EH 5.5kb plasmid as a template with a forward primer containing AatII and SmaI sites, and a reverse primer containing a PciI site, which causes a mutation of C to A at nucleotide +236 relative to the transcription start site. This nucleotide change is not present in the GOF constructs and did not affect expression, as evidenced by the D-450 deletion and GOF1 constructs having the same GUS activity pattern. To fuse the upstream region with the *β-glucuronidase* (*GUS*) reporter gene, the amplified fragment was digested with AatII and PciI and ligated to AatII- and NcoI-digested pGEM5GUS vector to make pGEM5GUSPCS336. pGEM5GUSPCS336 was digested with SmaI and NotI and the promoter/*GUS* fusion gene fragment was ligated to a pGV1500-derived plant transformation vector, pGV1501AN (12), to make D-4509. Fragments containing part of the *GA 20-oxidase* upstream region and part of the *GUS* gene were amplified by PCR using the D-4509 plasmid as a PCR template with forward primers containing an EcoRI restriction enzyme site. The PCR fragments were digested with EcoRI and PshAI and ligated into the EcoRI- and PshAI-digested D-4509 plasmid, generating the 5’ deletion constructs. The *GA 20-oxidase* fragment regions were sequenced. Primer sequences are listed in Table S3.

### Gain-of-Function (GOF) Constructs

Fragments containing part of the *GA 20-oxidase* upstream region were amplified by PCR using the D-4509 plasmid as a PCR template with primers containing either EcoRI or XmaI restriction sites. The amplified DNA fragments were digested with EcoRI and XmaI and ligated into pX46GV, a *β-glucuronidase* (*GUS*) reporter gene (40) vector that carries the *CaMV 35S* minimal promoter (41, 42). The *GA 20-oxidase* fragment regions were sequenced. Primer sequences are listed in Table S3.

### Site-Directed Mutagenesis Constructs

Predicted motifs were mutated according to the strategy we previously used in our laboratory (11). Transversional mutagenesis was used unless this process created a new predicted motif, in which case adenine substitution was used. For M1 and M2, Splicing by Overlap Extension (SOEing) PCR (43) was used to generate fragments containing the mutated motifs using the GOF2 plasmid as a template. The amplified fragments were digested with EcoRI and XmaI, and ligated to the EcoRI- and XmaI-digested GOF2 plasmid to create the mutated constructs. For M3 and M5-M8, the GOF2 plasmid was used as a PCR template with primers containing the desired mutation and EcoRI or XmaI restriction sites. The amplified fragments were digested with EcoRI and XmaI and ligated to the EcoRI- and XmaI-digested GOF2 plasmid. A fragment of M4 flanked by EcoRI and XmaI restriction sites was made by gene synthesis (Genewiz, South Plainfield, NJ). This fragment was digested with EcoRI and XmaI and ligated to EcoRI- and XmaI-digested GOF2, generating plasmid M4. The *GA 20-oxidase* fragment regions were sequenced to confirm that they contained the correct mutated bases. Primer sequences are listed in Table S3.

## Acknowledgements

We are grateful to present and past members of our laboratory for discussion and advice with this project, and especially Min Chen and Xiaomeng Wu for generating SRB and Common Bean embryo transcriptomes. We thank Professor John Harada for insightful comments on our suspensor *cis*-element research, as well as Professor Jeff Long for assistance with non-radioactive *in situ* hybridization. We dedicate this paper to the memory of our close friend and mentor, Professor Eric Davidson, who was a visionary in the field of developmental biology, a pioneer on the organization of eukaryotic gene regulatory networks, and provided us with perceptive advice on *cis*-regulatory module identification. This work was funded by a grant from the National Science Foundation Plant Genome Program.

## SUPPLEMENTARY FIGURES AND TABLES

**Fig. S1.**
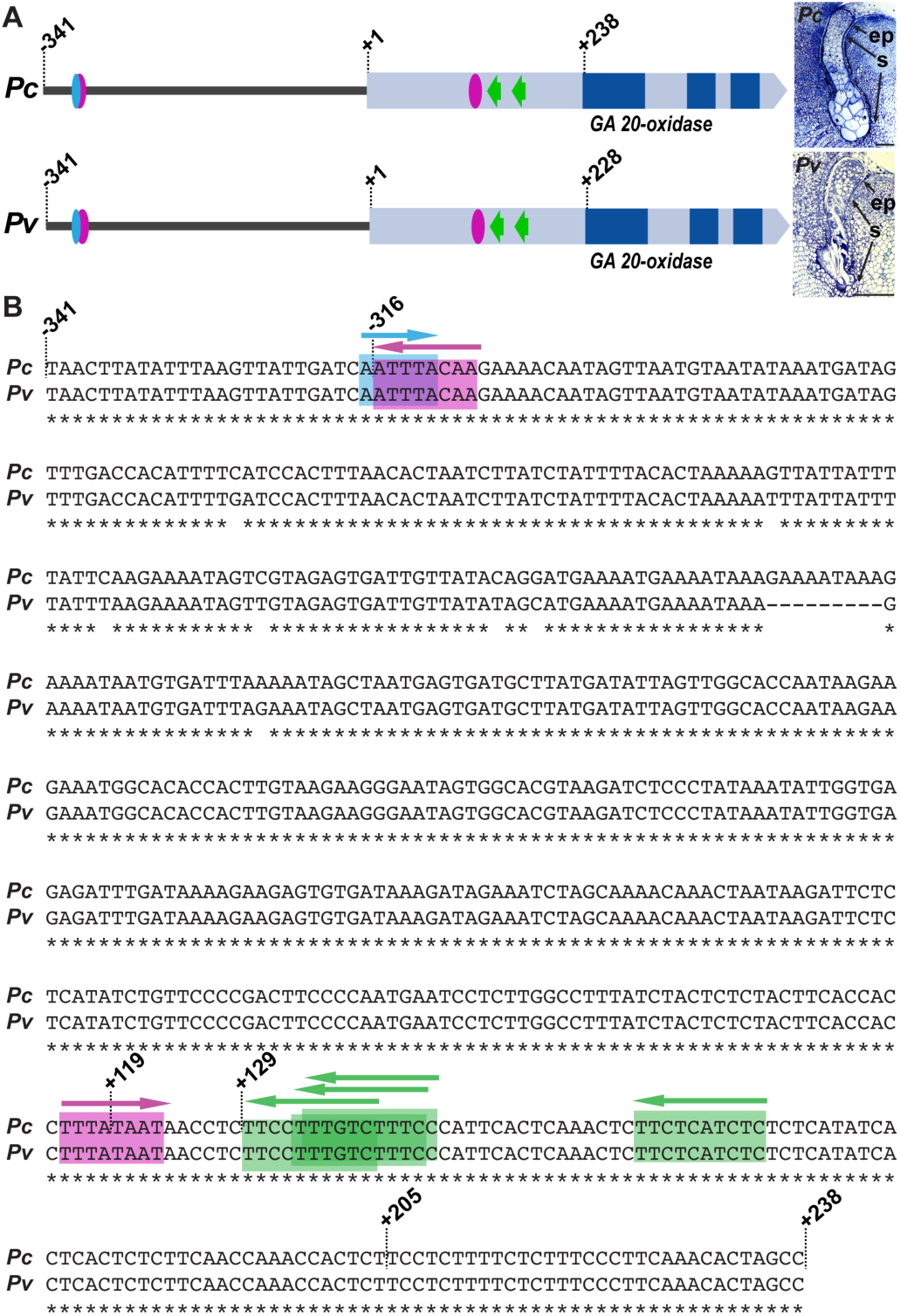
Conservation of suspensor *cis*-regulatory elements in the upstream region of *GA 20-oxidase* in SRB and Common Bean. (*A*) Conceptual representation of SRB and Common Bean *GA 20-oxidase* genes and location of functional suspensor *cis*-regulatory motifs. Green arrows, purple ovals and blue ovals indicate the 10-bp motif, Region 2 motif and Fifth motif, respectively. Dark blue boxes represent exons. Light blue boxes represent UTRs and introns. Numbers indicate nucleotide positions relative to the SRB transcription start site (+1). Suspensor images were reproduced from Fig. 1*B* and *D*. (*B*) Nucleotide sequence alignment of the SRB and Common Bean *GA 20-oxidase* upstream regions. Nucleotides conserved in SRB and Common Bean are indicated by asterisks. Green, purple and blue boxes indicate the 10-bp motif, Region 2 motif and Fifth motif, respectively. Arrows indicate the orientation of the motifs. ep, embryo proper; *Pc*, *Phaseolus coccineus*; *Pv*, *Phaseolus vulgaris*; s, suspensor. (Scale bar: 50 µm.)

**Table S1.**
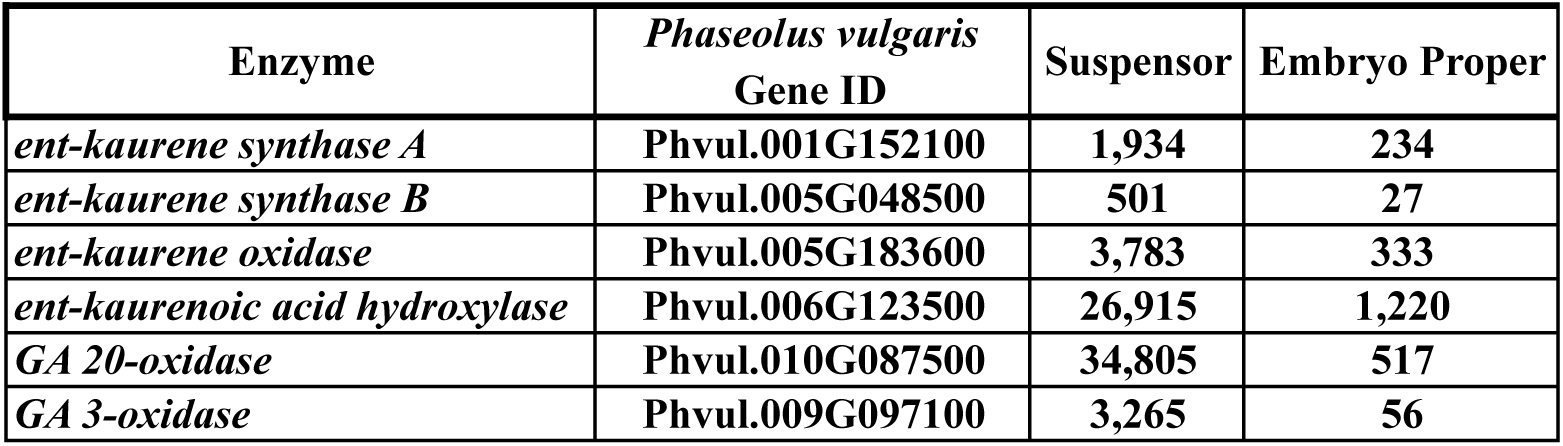
GA biosynthesis enzyme mRNA prevalence in the globular-stage suspensor and embryo proper. mRNA prevalence for each GA biosynthesis gene in SRB and Common Bean suspensor and embryo proper regions. RNA-Seq data were taken from GEO accession GSE57537. Numbers indicate average reads per kilobase per million (RPKM) of two biological replicates.

**Table S2.** Probes used for SRB *in situ* hybridization. Probes were generated from cDNA clones made from micro-dissected suspensor and embryo proper regions of globular-stage SRB embryos 6 DAP (12). SRB EST sequences were compared to the *Phaseolus vulgaris* protein database (http://phytozome.jgi.doe.gov) using BLASTX, and the top *Phaseolus vulgaris* gene ID is displayed for each SRB EST (44).

| Enzyme | GenBank<br>Accession<br>Number | SRB<br>EST<br>Length | *Phaseolus vulgaris*<br>Gene ID |
| --- | --- | --- | --- |
| *ent-kaurene synthase A* | CA914587 | 415 | Phvul.001G152100 |
| *ent-kaurene synthase B* | CA899647 | 526 | Phvul.005G048500 |
| *ent-kaurene oxidase* | CA914551 | 466 | Phvul.005G183600 |
| *ent-kaurenoic acid hydroxylase* | CA914608 | 257 | Phvul.006G123500 |
| *GA 20-oxidase* | CA914823 | 582 | Phvul.010G087500 |
| *GA 3-oxidase* | CA915026 | 733 | Phvul.009G097100 |

**Table S3.**
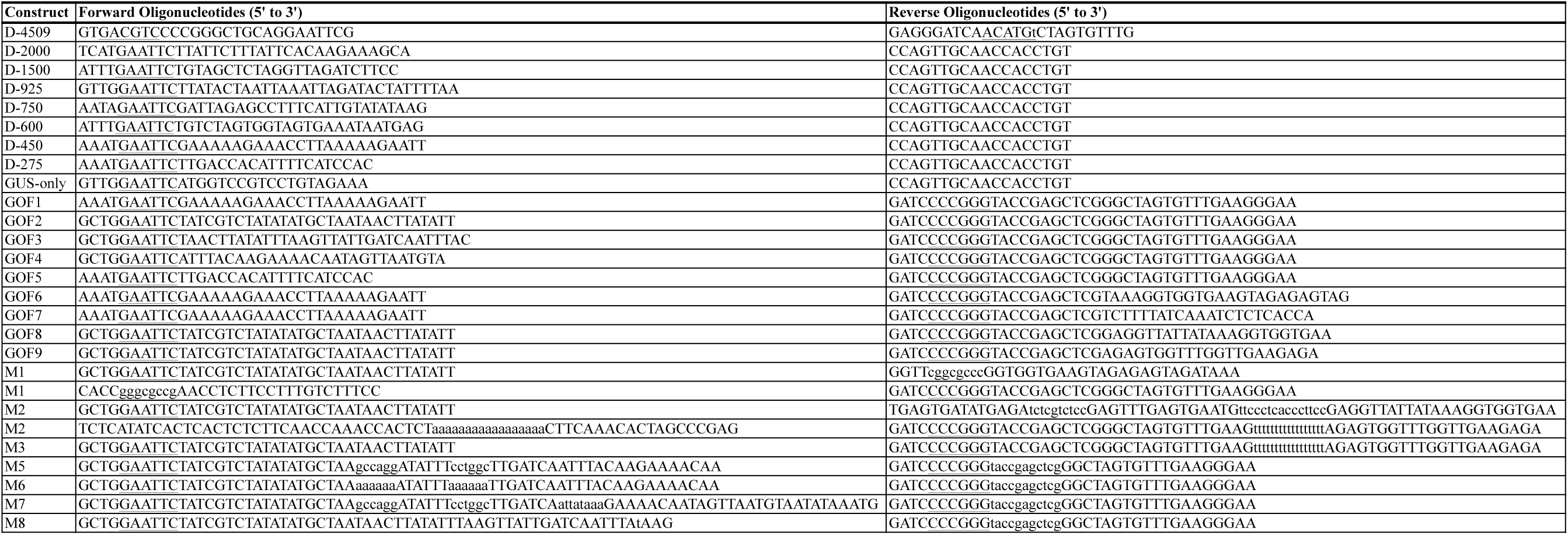
Oligonucleotide sequences for generating constructs. Underlined nucleotides are incorporated restriction sites. Lower case nucleotides are mutated relative to the *GA 20-oxidase* upstream sequence.

**Author contributions:** K.F.H., A.Q.B., T.K. and R.B.G. designed research; K.F.H., A.Q.B. and T.K. performed research; K.F.H., A.Q.B., T.K. and R.B.G. analyzed data; and K.F.H. and R.B.G. wrote the paper.

